# Evaluating the contribution of genome 3D folding to variation in human height using machine learning

**DOI:** 10.1101/2025.09.09.675195

**Authors:** Wanjun Gu, Erin N. Gilbertson, Sergio E. Baranzini, Rany S. Salem, John A. Capra

## Abstract

Genome-wide association studies (GWAS) have identified thousands of variants associated with complex traits, yet the majority lie in noncoding regions, making it difficult to determine their functional impact. Alterations to the three-dimensional (3D) spatial interactions among gene regulatory elements are increasingly recognized as a mechanism by which genetic variants influence gene expression. However, experimentally evaluating whether variants disrupt 3D-genome structure is not feasible at GWAS scale. To address this, we developed a computational framework that integrates GWAS summary statistics with predictions from the Akita sequence-based deep learning model of 3D chromatin contacts. We applied the framework to 9,917 genomic regions associated with human height, assessing both individual variants and haplotypes for their predicted impact on 3D genome architecture. Only a small fraction of height-associated haplotypes had substantial predicted disruption of 3D folding (17 regions, 0.17%, exceeded a disruption score of 0.1). Considering all common variants in a haplotype together generally produced greater perturbations than individual variants, but several highly divergent regions were driven by single variants. We highlight a variant that disrupts the binding motif at a confirmed CTCF binding site and is predicted to modify 3D genome contacts with the *LCOR* promoter, suggesting that 3D-genome-mediated disruption of gene regulation underlies the association with height. This work presents a scalable and interpretable strategy for integrating 3D genome modeling with GWAS, enabling investigation of this important regulatory mechanism in the connection of non-coding genetic variation to complex traits.

## Introduction

The mechanisms through which genetic variation contributes to phenotypic diversity remain unresolved for most complex traits^1–3^. Complex traits, including anthropometric traits, such as body height^4,5^ and weight^6–8^, and heritable diseases, such as diabetes^9–11^, and psychiatric disorders^12,13^, exhibit polygenic architectures involving numerous genetic loci across the genome, each potentially influencing the phenotype through distinct molecular mechanisms and biological pathways. However, small effect sizes and limited statistical power pose significant challenges to isolating the precise roles of individual variants^14–16^. Additionally, ancestry-specific linkage disequilibrium (LD) patterns can further complicate interpretation, as association signals detected in GWAS may not reflect the causal variant itself but rather a nearby variant in strong LD. These correlated variants can mask the true biological mechanism underlying the association. LD structure also differs across populations due to demographic factors such as bottlenecks and admixture, making trans-ancestry fine-mapping particularly difficult^17^. However, new genomic technologies are enabling the quantification of molecular and cellular attributes relevant to gene regulation and phenotypic variation^18^.

The three-dimensional (3D) genome architecture, defined by the frequency of physical contacts between distal genomic regions, plays a critical role in gene regulation by enabling interactions between regulatory elements such as enhancers and promoters^19,20^. Data from consortia such as the 4D Nucleome Project^21^ have provided valuable insights into the organization of chromatin and its influence on gene expression. Studies leveraging the 3D-genome structure data have shown that the spatial proximity of regulatory elements, such as enhancers and promoters, can be predictive of transcriptional activity, and that disruptions in 3D genome organization, such as altered topologically associating domains (TADs) or loop formations, can lead to cell-type-specific dysregulated gene expression and different disease outcomes^20,22–26^. Disruptions to 3D-genome architecture have been linked to both rare and complex diseases by altering long-range regulatory interactions and promoting genomic rearrangements^27^. These disruptions can interfere with long-range regulatory interactions, leading to gene misexpression, enhancer hijacking, or ectopic chromatin interactions^28–30^. For instance, structural variants that alter TAD boundaries have been shown to cause limb malformations^31^, cancers^32^, and neurodevelopmental disorders^32,33^. However, the full extent of 3D-genome variation across genetically diverse populations is still not fully understood, due to the prohibitive cost of quantifying 3D genome variation in large cohorts.

Recent advances in machine learning allow prediction of 3D chromatin contacts from DNA sequences, complementing the costly and time-intensive Hi-C experiments. Akita, a convolutional neural network, was trained on high-resolution Hi-C and Micro-C data from five human cell types and can predict contact frequency maps from 1 Mb DNA sequence windows at ∼2 kb resolution^34^. Akita has demonstrated robust performance, with the median of test-set Spearman correlations exceeding 0.55 between predicted and experimental contact maps on held-out chromosomes, approaching the concordance observed between experimental replicates (often ranging between 0.6 to 0.7). Notably, Akita captures locus-specific folding patterns, learns predictive sequence features, and has been widely used in eQTL interpretation^35^ and structural variant validation^36^. This computational tool also opens the door for population-level analyses of how genetic variants of interest affect 3D-genome architecture^37^. However, despite these advances, the potential role of 3D genome folding disruption on complex trait variation at the population scale remains largely unexplored.

To investigate the relevance of 3D genome folding to complex trait variation, we focused on human height, a well-characterized, highly heritable trait. A recent genome-wide association study (GWAS) from the GIANT consortium involving over five million individuals from diverse ancestries has provided a comprehensive map of its genetic architecture^5^. Height is particularly well-suited for exploring regulatory mechanisms, including those involving 3D genome organization, due to its robust replication across cohorts, minimal environmental confounding, and highly polygenic nature.

We used sequence-based machine learning to predict 3D-genome contact maps for height-associated genetic loci^5^. We leveraged large-scale phased genome data from the TOPMed consortium^38^ to compute haplotypes and investigate the impacts of the height-associated variants they carry on 3D genome contacts. While 3D genome folding disruption is only rarely predicted to contribute to height variation, we nominate several height-associated variants that likely modify 3D genome contacts containing known regulatory elements. Our results offer new insights into the complex relationship between genome architecture and height-associated loci. This study demonstrates a novel framework to functionally interrogate the potential effects of GWAS-significant variants on a crucial regulatory mechanism through sequence-based machine learning.

## Methods

This study explores the role of three-dimensional (3D) genome disruption as a mechanism underlying height-associated genetic loci. We leverage multiple publicly available genomic resources and machine-learning-based in-silico predictions of genome folding. The analysis pipeline (Figure 1) to examine the effect of 3D-genome disruption consists of genome-wide association study (GWAS) data processing, GWAS significant variant identification, haplotype inference, haplotype-aware in-silico mutagenesis, and machine learning predictions of 3D-genome disturbance.

**Figure 1.**
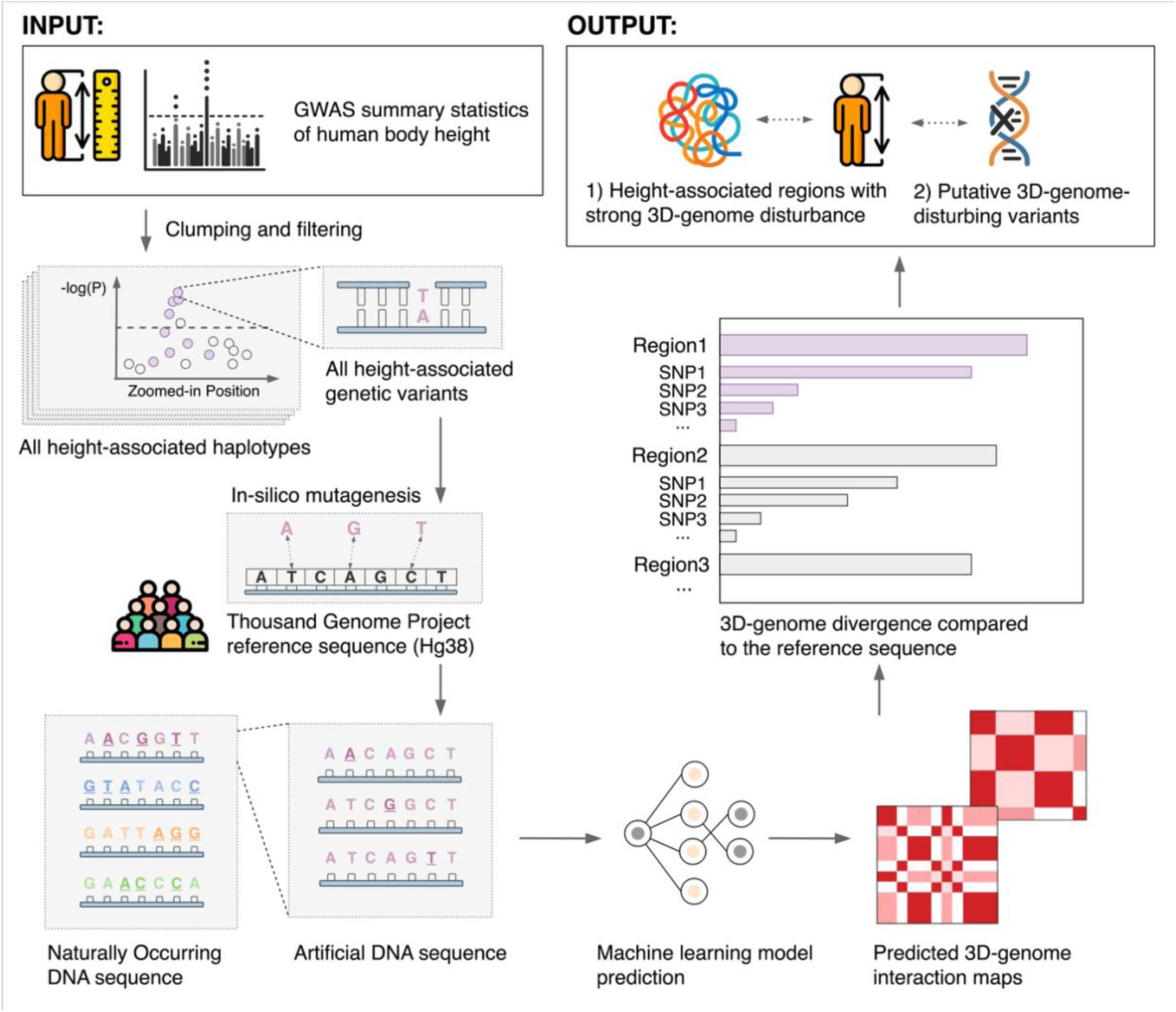
Overview of the analysis pipeline for assessing the impact of 3D-genome divergence in height-associated genomic loci. The analysis begins with input from GWAS summary statistics for human body height, from which height-associated loci are identified using clumping and filtering based on linkage disequilibrium and statistical significance thresholds. All variants within associated loci are collected and grouped into haplotypes. Using the 1000 Genomes Project reference panel of build GRCh38/Hg38, both naturally occurring haplotypes and single-variant-edited sequences are generated through in silico mutagenesis. These sequences are input into a DNA-sequence-based machine learning model (Akita) to predict 3D chromatin contact maps. Divergence is quantified by comparing predicted maps to the reference sequence using Spearman’s correlation across the contact maps. Regions are ranked based on the magnitude of 3D-genome divergence, and putative 3D-genome disturbing variants are identified for the top height-associated regions with strong 3D architectural disruption.

### GWAS data preprocessing and significant locus identification

Summary statistics were downloaded from the recent GIANT GWAS on standing height^5^. Genetic loci with suggestive associations (p ≤ 1 × 10^−5^) with body height were identified through clumping and filtering of GWAS signals to select independent variants based on linkage disequilibrium (LD). The suggestive significance threshold (p ≤ 1 × 10^−5^)^39,40^ was used to inclusively identify candidate associations for secondary analyses, capturing a broad set of loci that may not reach genome-wide significance but could still represent biologically relevant and potentially functional variants. Both clumping and filtering were conducted using PLINK2^41^. The LD threshold for clumping was an r^2^ of 0.5 and single nucleotide polymorphisms (SNPs) within 250 kb from the lead SNP were considered for clumping. These commonly used^1,42,43^ parameters strike a balance between collapsing redundant signals and preserving distinct association signals, ensuring that nearby variants in moderate linkage disequilibrium are grouped without over-merging biologically independent loci. The GRCh38/hg38 genome assembly was used throughout the analysis to ensure consistency across datasets and accurate mapping of variants.

### Haplotype imputation using TOPMed sequencing data reference

To perform haplotype-aware analyses, the NHLBI Trans-Omics for Precision Medicine (TOPMed) whole-genome sequencing datasets were used as a haplotype reference^38^. The curated dataset, comprising ∼50,000 individuals from diverse ancestries, including European, African, East Asian, South Asian, and admixed/non-admixed American populations, allowed haplotypes present within the significant height-associated loci to be imputed and phased variants within the credible sets to be included in the analysis.

### In-silico mutagenesis and sequence generation

Using the human reference sequence (hg38) and the 1000 Genomes Project^44^, we generated the sequences of all naturally occurring haplotypes with a count ≥ 30 in TOPMed for all height-associated loci to ensure adequate representation and reliability in downstream analyses, while reducing the burden of evaluating rare haplotypes^45^. Furthermore, within each height-associated locus, single SNPs were edited into the reference sequence to examine the consequences of single variants. This produced two sequence sets for *in-silico* mutagenesis: 1) naturally occurring DNA sequences, denoted as haplotypic sequences, and 2) artificial DNA sequences, denoted as single-SNP altered sequences.

### Prediction of 3D-genome disturbance using machine learning

Akita, a convolutional neural network capable of predicting 3D genome structure from DNA sequence^34^ was used to quantify the impact of both natural and artificial sequences on genome folding. For this study, we used the HFF model, which was trained on high-resolution Hi-C and Micro-C data from the human foreskin fibroblast cells. This cell type was selected because it has the deepest sequencing coverage and the highest predictive performance among the cell types used in Akita’s training. The model takes as input a ∼1 megabase (Mb, 2^20^ bp) DNA sequence, and outputs a predicted 3D contact frequency map at ∼2 kilobase (2^11^ bp) resolution across the entire input region, resulting in a matrix of predicted interactions between all ∼2 kb segments within the 1 Mb window. The contact frequency maps of each sequence derived from the reference (naturally occurring and variant-induced) were then compared to one another and the contact frequency map of the reference sequence. We computed Spearman’s rank correlation coefficient (ρ) between each edited sequence and the reference sequence, and following previous work, we refer to *D* = 1 − *ρ* as the 3D genome divergence^37^. Thus, a larger 3D genome divergence indicates a larger difference between the contact maps.

### Ranking and annotation of 3D-genome-diverged regions

We ranked the height-associated loci based on the 3D genome divergence compared to the reference sequence. We used quantile-quantile regression to determine loci whose 3D-genome divergence is significantly larger than expected. The 3d-genome divergence was defined as *D*_*i*_ for each locus *i*. A base-10 logarithmic transformation was applied to better visualize the distribution and stabilize variance. Log-transformed *D*_*i*_ values were assumed to be approximately normally distributed. Upon sorting the observed *D*_*i*_ in ascending order, the theoretical quantile of *D*_*i*_ can be computed from a standard normal distribution using:

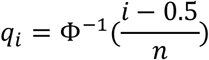

where Φ^−1^ is the inverse cumulative distribution function (CDF) of the standard normal distribution, and *n* is the number of loci. The ranks of observed *D*_*i*_ that were significantly positively deviated from the theoretical quantiles *q*_*i*_ are candidate 3D-genome disturbing loci. A subset of the regions with highest predicted disturbance were selected for further analysis. Functional annotations were assigned to these regions of interest based on nearby genes and available functional genomics data.

### Identification of putative functional variants within regions of interest

To identify putative functional variants within height-associated regions, the 3D-genome divergence of both the natural and artificial sequence from the reference was compared. The SNP contribution to 3D-genome disturbance was quantified based on the ratio between the disturbance of the SNP and the maximum disturbance introduced by a naturally occurring haplotype.

### Enrichment quantification of loci showing 3D-genome divergence

We used linkage disequilibrium score (LD score) regression^46,47^ to determine whether 3D genome divergent loci associated with height were either: 1) enriched for heritability among all height-associated loci, or 2) enriched for heritability among all regions associated with 3D-genome divergent loci. The clumped and filtered summary statistics for body height were harmonized to the HapMap3 reference panel^48^ to ensure consistency. SNPs within loci predicted to have 3D genome divergence were identified using BEDTools^49^ by intersecting the genomic coordinates of the GWAS summary statistics and the 3D genome data. Partitioned LD scores were calculated using reference data from the 1000 Genomes Project (Phase 3, European ancestry).

## Results

### A small number of height-associated loci have evidence of 3D genome contact divergence

We analyzed haplotypes and SNPs from 9,917 height-associated genomic regions to predict their impact on 3D-genome architecture. The distribution of 3D-genome disturbance for both the naturally occurring as well as the artificial sequences follow an expected distribution with the inflation index lambda approximately equal to one (λ=1.0002 for naturally occurring; λ=0.9998 for artificial sequences), suggesting no inflation or deflation of the 3D-genome disturbance prediction (Figure 2A). Only a small fraction of these loci exhibited substantial 3D-genome disturbance. We defined notable divergence as D ≥ 0.01 (∼99^th^ percentile), and extreme divergence as D ≥ 0.1. These thresholds correspond to effect sizes previously shown to correlate with regulatory impact in models of in silico chromatin contact prediction and were further supported by empirical inspection of the divergence distribution^34^. Specifically, 107 regions (1.08%) demonstrated notable divergence, and 17 regions (0.17%) displayed extreme divergence in 3D architecture (Table1; Figure 2B). The top 17 regions of 3D-genome disturbance are shown in Table 1 with their respective genes and ancestry-specific GWAS summary statistics.

**Figure 2.**
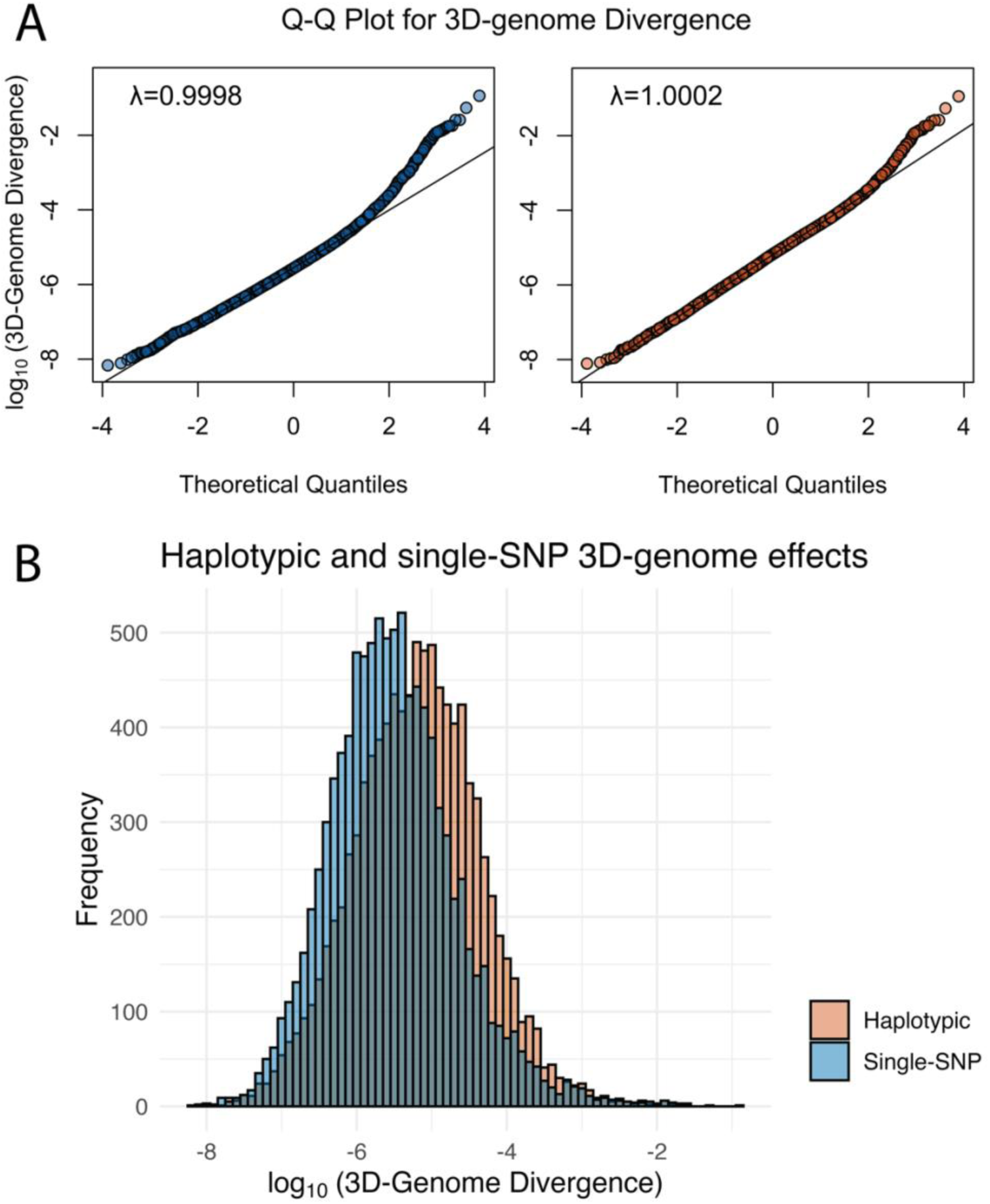
Haplotypic and single-SNP 3D-genome effects. (A) Q-Q plots show the alignment of observed log₁₀ 3D-genome divergence values against theoretical quantiles under a normal distribution. Both plots (left: artificial single-SNP-edited sequences, λ=0.9998; right: naturally occurring haplotype sequences, λ=1.0002) demonstrate that the observed values closely follow the expected distribution, with slight deviations at the tails. (B) The histogram compares the distribution of log₁₀-transformed 3D-genome divergence values between naturally occurring haplotype sequences (orange) and artificial single-SNP-edited sequences (blue). Haplotypes display higher divergence scores and a broader distribution, indicating that haplotype sequences produce larger 3D-genome divergence on average than individual SNPs. The higher frequency of larger divergence values for haplotypes suggests cumulative effects of multiple variants within the same haplotype.

**Table 1.** Summary of height-associated SNPs with the largest inferred 3D-genome divergence. Table columns: 1) SNP: The genomic location and alleles for the top height-associated SNP at a locus given as Chromosome:Position:Reference allele:Alternative allele. All positions are reported in the hg38 genome assembly. 2) rs#: The rs number of the SNP. 3) Gene: The overlapping gene for each SNP. For SNPs in intergenic regions, no genes were listed. 4) Divergence: The maximum 3D-genome divergence (1 – Spearman’s correlation) induced by naturally occurring height-associated haplotypes. All loci with divergence greater than 0.01 are reported. 5) GWAS Ancestry: The ancestry (EUR: European, EAS: East Asian) of the GWAS from which the significant association with body height was identified. 6) P-Value: The P-value of the SNP’s association with height. 7) Effect Size: The effect size of the association, representing the magnitude of the SNP’s impact on height. The unit of the effect size is expressed in terms of standard deviations per minor allele for SNP effect estimates. 8) EAF: The allele frequency of the effect allele across the studied population.

| SNP | rs# | Gene | 3D Divergence | GWAS Ancestry | P-Value | Effect Size | EAF |
| --- | --- | --- | --- | --- | --- | --- | --- |
| chr10:96789820:T:C | rs7477274 | *LCOR* | 0.113 | EUR | 6.30E-14 | 0.014 | 0.09 |
| chr3:46258274:A:G | rs7652290 | *CCR3* | 0.054 | EUR | 2.44E-09 | 0.007 | 0.32 |
| chr12:104877307:G:T | rs7299019 | *SLC41A2* | 0.026 | EAS | 8.75E-10 | 0.021 | 0.25 |
| chr9:34011393:G:A | rs12555291 | *UBAP2* | 0.026 | EUR | 4.56E-08 | -0.006 | 0.34 |
| chr4:122914501:C:A | rs12513181 | *NUDT6* | 0.024 | EUR | 4.38E-58 | -0.019 | 0.74 |
| chr9:34382728:A:G | rs3808876 | *SPMIP6* | 0.019 | EUR | 3.22E-23 | 0.011 | 0.57 |
| chr6:104897150:T:C | rs17065381 | *LIN28B-AS1* | 0.018 | EUR | 8.54E-54 | 0.034 | 0.06 |
| chr9:106150630:A:G | rs1516882 | *LINC01505* | 0.018 | EUR | 1.27E-99 | -0.024 | 0.32 |
| chr14:65053528:T:C | rs2296316 | *FNTB* | 0.015 | EUR | 6.63E-95 | -0.022 | 0.46 |
| chr5:78493049:G:C | rs16875509 | *LHFPL2* | 0.014 | EAS | 8.27E-10 | 0.02 | 0.30 |
| chr4:45927155:A:G | rs6817621 |  | 0.014 | EUR | 3.06E-10 | -0.007 | 0.47 |
| chr14:24058463:G:A | rs7146599 | *CARMIL3* | 0.013 | EUR | 2.99E-18 | -0.01 | 0.50 |
| chr5:42019008:A:G | rs2011105 |  | 0.013 | EUR | 1.55E-19 | -0.02 | 0.06 |
| chr19:37912459:T:C | rs8105305 | *SIPA1L3* | 0.012 | EUR | 2.49E-33 | -0.013 | 0.58 |
| chr11:31399033:G:A | rs17632324 | *DNAJC24* | 0.012 | EUR | 1.97E-13 | -0.008 | 0.36 |
| chr5:72514419:G:A | rs246562 |  | 0.012 | EUR | 6.29E-22 | -0.013 | 0.19 |
| chr12:5428946:G:A | rs1861835 |  | 0.011 | EUR | 5.21E-14 | -0.013 | 0.09 |

### Haplotype effects on 3D divergence are larger than single-SNP effects

Next, we evaluated whether considering full haplotypes vs. individual SNPs produced different distributions of predicted 3D divergence. The effects of haplotypes on 3D-genome folding were found to be generally larger than those observed from individual SNPs. The average divergence caused by haplotypes was 9.37 × 10^−5^, significantly higher (p < 2.2 × 10^−16^) than 7.85 × 10^−7^ for single-SNP variants (Figure 2B). This observation indicates that, as expected, haplotype sequences with multiple variants are predicted to produce greater 3D divergence on average than individual SNPs.

### 3D genome divergence and GWAS effect size are not correlated

To investigate whether genetic loci with greater 3D genome divergence have larger effects on height, we analyzed all height-associated haplotypes by correlating the log-transformed magnitude of the maximum predicted 3D genome divergence of the height-associated haplotypes with the magnitude of their effect size on height. The analysis revealed no significant correlation between 3D genome divergence and effect size among all height-associated haplotypes (Supplementary figure S1A, β = 0.44, p > 0.527) or among haplotypes in the top 1% for 3D genome divergence (Supplementary figure S1B, β = −2.73, p > 0.423). These findings are consistent with the limited number of height-associated loci exhibiting large 3D genome divergence.

### Regions with 3D genome divergence are not enriched overall for heritability of height

Next, we performed partitioned heritability analysis using LD Score Regression^47,50^ to test whether body height heritability is enriched in previously dentified^37^ sets of 1-Mb genomic regions predicted by the Akita model to alter 3D genome folding. These 3D-genome divergent results were evaluated separately in the European (EUR), African (AFR), Admixed American (AMR), and East Asian (EAS) cohorts, as well as in the meta-analyzed GWAS summary statistics across all ancestries.

In each analysis, the 3D-genome-altering regions comprised approximately 6.6% of SNPs. Across cohorts, the proportion of height heritability attributed to these regions ranged from 6.7% to 9.5%, closely mirroring the SNP proportion. Correspondingly, enrichment estimates were close to one (1.43 in EUR, 1.26 in AFR, 1.32 in AMR, 1.02 in EAS, and 1.52 in the meta-analysis), with wide confidence intervals and non-significant p-values (Supplementary table S1). Taken together, these results indicate that there is not significant enrichment of height heritability within the predicted 3D-genome-altering regions, suggesting that the disruption of 3D-genome architecture is likely not a major factor driving variation in height.

### Several height-associated loci 3D divergent loci show predicted haplotype effects

To evaluate whether the height-associated loci with the largest 3D divergence are likely the result of a single variant or a combination of variants on the haplotype, we computed the percentage of the maximum 3D divergence observed for a haplotype that could be explained by the most disruptive single SNP.

Most highly divergent regions are dominated by the effect of a single SNP (Figure 3). However, we identified three exceptions where the maximum divergence explained by a single SNP was less than 60%. These were the chr4:122914501 locus within the *NUDT6* gene, the chr7:27870446 locus within the *JAZF1* gene, and the chr12:57852573 locus near the *CTDSP2* gene (Figure 3). In these cases, the top contributing variant accounted for less than 60% of the total disturbance, with multiple other variants in the haplotype contributing variation. These results suggest that for a small number of haplotypes with multiple variants may influence trait-associated 3D-genome architecture in combination.

**Figure 3.**
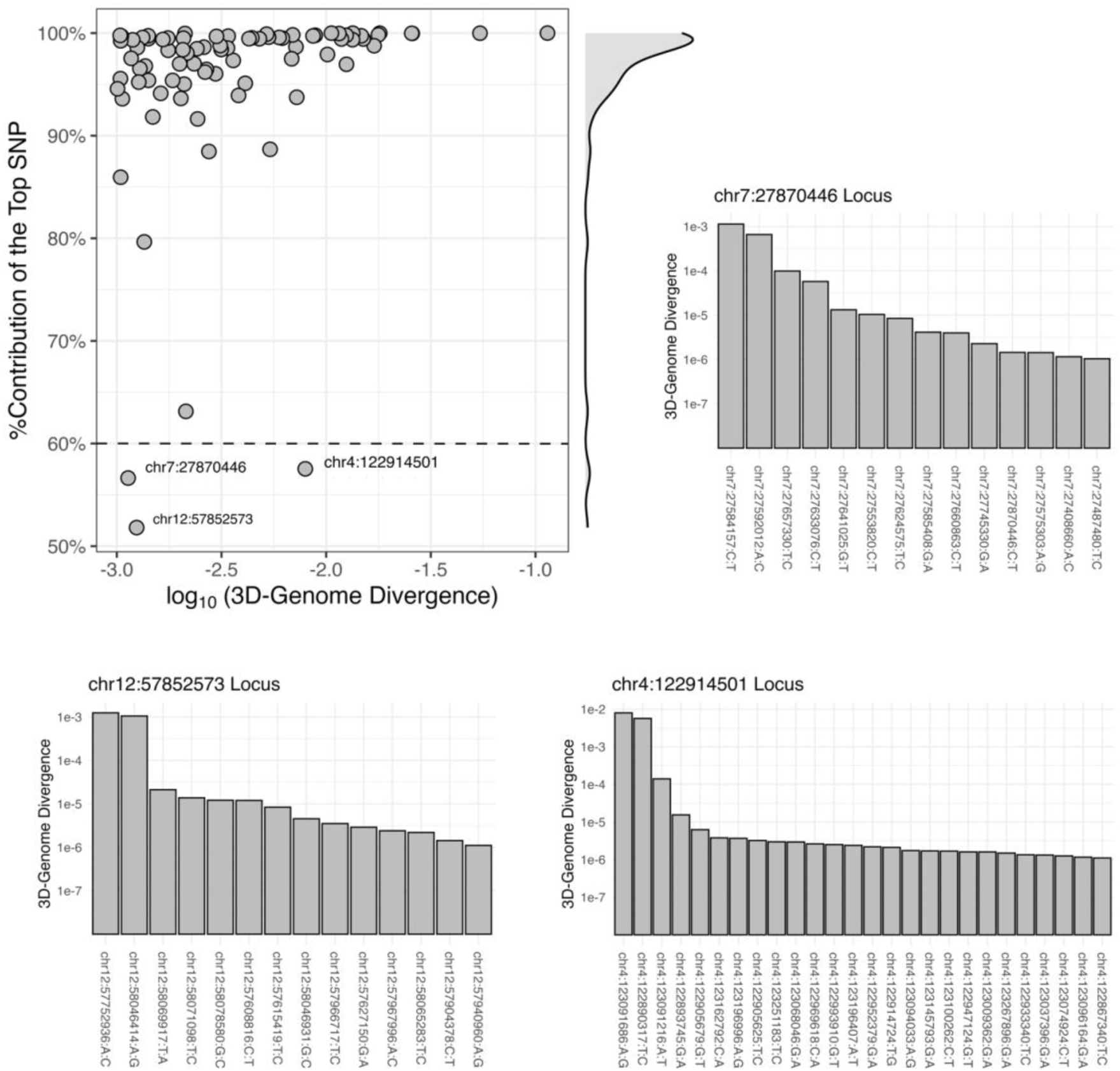
Contribution of individual SNPs to 3D-genome divergence across height-associated loci. The scatter plot (top left) illustrates the relationship between the log₁₀-transformed 3D-genome divergence and the percentage contribution of the top SNP within each height-associated region. For most regions a single SNP accounts for nearly all the predicted 3D-genome disturbance, with many clustered near 100%. However, several loci exhibit more complex haplotypic effects with the top SNP contributing less than 60% to the total divergence predicted for the haplotype (dashed line). Three height-associated loci, chr4:122914501 (*NUDT6*), chr7:27870446 (*JAZF1*), and chr12:57852573 (*CTDSP2*), are highlighted, each displaying significant cumulative contributions from multiple variants. The bar plots (right and bottom) provide detailed breakdowns of SNP-level contributions to 3D-genome divergence within the three loci. In these cases, the top contributing SNP explains only a portion of the disturbance, with several additional variants contributing to the overall 3D-genome divergence. This pattern suggests haplotypic effects, where multiple variants collectively influence chromatin architecture.

### The top height-associated locus with evidence of 3D divergence disrupts a CTCF binding site in the promoter of LCOR

The *LCOR* gene overlaps, chr10:96789820 (T>C), a strongly height-associated SNP (p = 2.44 × 10^−9^), with the highest predicted 3D-genome divergence among all loci tested (0.113). LCOR functions as a transcriptional repressor involved in chromatin remodeling^51^, suggesting that modification of the regulation of this gene could interfere with critical regulatory processes. Comparing the naturally occurring haplotypes with the reference (T) and alternative allele (C) demonstrated significant divergence in 3D genome contacts with the *LCOR* promoter (Figure 4A). The chr10:96789820 variant accounts for over 99.9% of the observed 3D-genome divergence in this region (Figure 4B). We found that the T>C change at this locus disrupts the sequence motif for the binding of the CTCF protein, a crucial regulator of genome 3D contact patterns. Based on ChIP-seq data from the ENCODE project^52^, the binding of CTCF at this locus is strongly supported, with CTCF being detected frequently across diverse biosamples and tissue types (Figure 4D).

**Figure 4.**
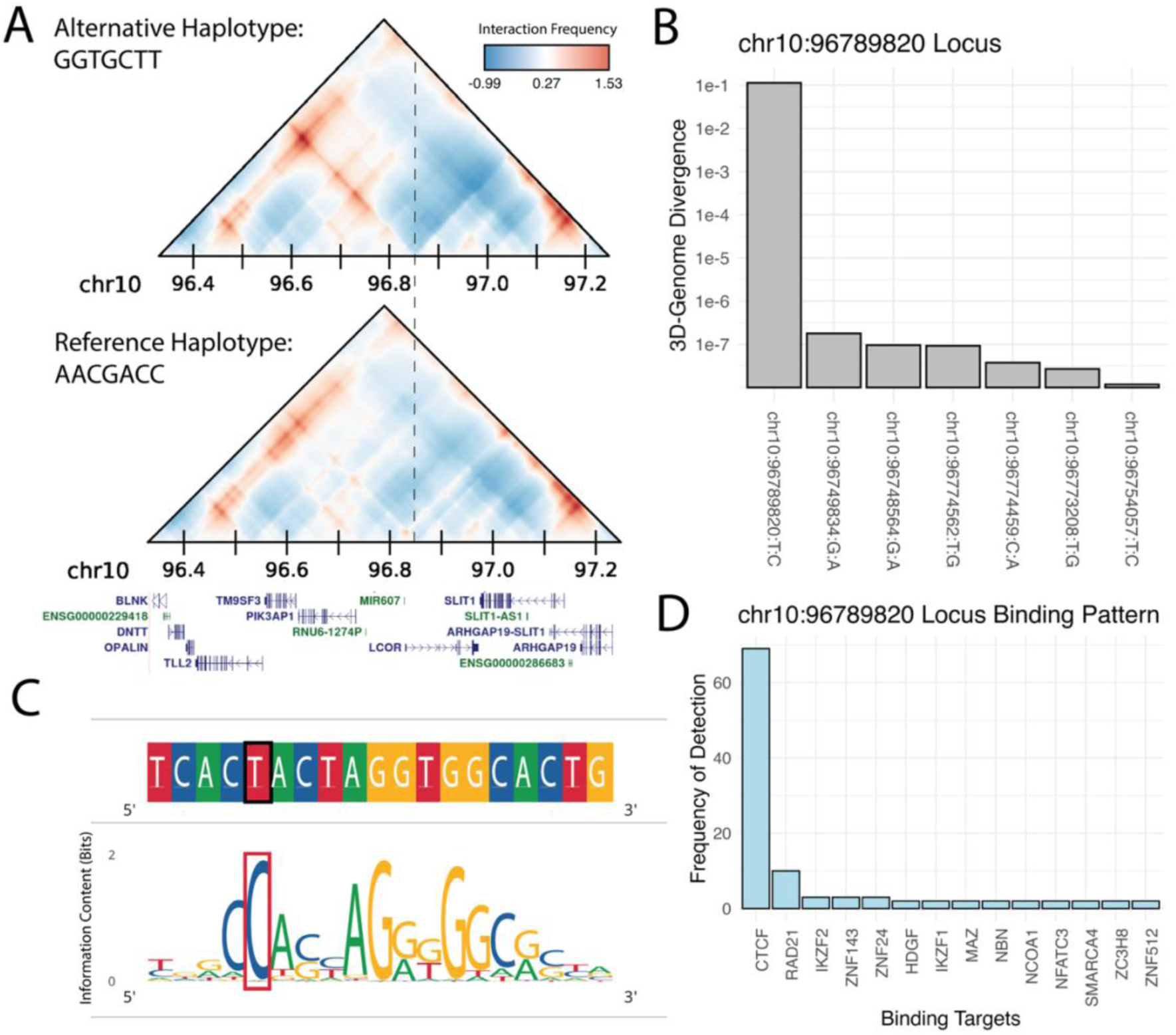
A height-associated variant disrupts a CTCF motif and inferred 3D contacts at the *LCOR* promoter. (A) Predicted interaction maps of the alternative haplotype (GGTGCTT) and the reference haplotype (AACGACC). The heatmaps visualize predicted contact frequencies within the 96.4–97.2 Mb region of chromosome 10. Stronger red values indicate higher relative contact frequencies, while blue represents fewer contacts than expected. The alternative haplotype shows a clear shift in chromatin contacts with regions upstream of the *LCOR* promoter (dashed line) compared to the reference haplotype, suggesting that this variant alters 3D-genome contacts with regulatory functions. (B) Bar plot of individual SNP 3D-genome divergence at the chr10:96789820 locus. The chr10:96789820T>C variant alone accounts for the largest share of divergence, with additional SNPs contributing smaller effects. (C) Sequence motif analysis around the chr10:96789820 variant, showing the reference and alternative alleles. The red box highlights the disruption of a CTCF binding motif, a key factor in maintaining chromatin structure, caused by the T>C substitution. (D) Bar plot summarizing protein binding at the chr10:96789820 locus based on ChIP-seq data from the ENCODE project. CTCF binding is detected across more than 60 contexts, reinforcing the importance of this site in genome organization. This result suggests that the T>C substitution at this locus likely alters chromatin architecture by disrupting CTCF binding, potentially affecting gene regulation of *LCOR*.

## Discussion

This study provides insights into the relationship between 3D-genome folding and genetic loci associated with body height. We have two main conclusions. First, we identify a small number (∼15) height-associated loci with evidence that disruption of genome 3D contacts is the mechanism by which they influence variation in this trait. Second, beyond these examples, our results suggest little contribution of 3D genome contact disruption to variation in height, suggesting that the functional mechanism of action of the majority of height associated variants does not involve 3D genome contact disruption.

For most of the height-associated loci with 3D divergence, a single variant was predicted as the primarily driver of the change, suggesting that single-variant changes remain the dominant model for disrupting 3D-genome folding. However, three loci stood out as exceptions, *NUDT6*, *JAZF1*, and *CTDSP2*. Multiple variants within haplotypes collectively were predicted to cause 3D-genome divergence.

These results highlight the importance of haplotype-resolved analysis, as the large effects on 3D genome contacts were only detected when considering the combined effects of multiple variants rather than individual SNPs. The *NUDT6* gene, involved in mRNA de-capping and the regulation of cell proliferation, plays a role in pathways critical to growth and development, making it a biologically plausible candidate for height regulation^53,54^. Disruptions in chromatin structure at this locus may alter gene expression in ways that affect cellular growth dynamics, influencing final body height. Similarly, the *JAZF1* locus is involved in metabolism, insulin sensitivity, and growth regulation^55,56^. Here, 3D-genome disruption may impact enhancer-promoter interactions, modulating JAZF1 expression to affect growth processes alongside metabolic pathways. Finally, the *CTDSP2* locus, involved in transcriptional regulation via the dephosphorylation of RNA polymerase II, suggests that changes in 3D-genome architecture could disrupt developmental transcriptional programs, contributing to height variability^57^. These predictions indicate that cumulative haplotypic effects can reshape chromatin interactions, suggesting that variants at these sites may require finely coordinated regulation to maintain proper genome folding and gene expression. The identification of these loci underscores the need for haplotype-aware approaches in the study of the mechanisms underlying GWAS associations in our efforts to understand the regulatory mechanisms that influence variation in complex traits such as height.

Several factors likely explain why predicted 3D-genome divergence is not correlated with GWAS effect size in this study. First, the vast majority of genetic variants influencing height appear to act through regulatory mechanisms that do not require substantial alterations to local 3D chromatin conformation, such as modulating enhancer strength^58^, splicing efficiency^59^, or mRNA stability^60^. Second, modest but biologically meaningful changes in chromatin organization may not be fully captured by global divergence metrics like *D* = 1 − *ρ*, which summarize genome-wide contact differences rather than focusing on specific regulatory loops. It is also possible the functionally relevant contact changes happen at scales smaller than the resolution of the 2 kb bins used for prediction. Finally, effects on 3D genome contacts could be highly cell-type specific, and thus difficult to capture. Together, these possibilities underscore the complexity of the relationship between chromatin architecture and gene regulation and suggest that 3D-genome perturbation may represent a complementary, rather than predominant, mechanism underlying most height-associated genetic effects.

The sequence-based machine learning approach described here has several attributes that complement traditional ancestry-based fine-mapping methods for identifying causal variants from GWAS. Unlike traditional approaches, which rely on population diversity to identify functional variants and struggle in regions with strong linkage disequilibrium, this method can directly analyze variants in any combinations regardless of local LD patterns. By predicting 3D-genome folding from DNA sequences, the analysis pipeline of this study provides a powerful in-silico perspective for identifying putative functional variants, reducing the need for time-consuming gene-editing experiments. Importantly, the present study applies this framework to test a defined mechanistic hypothesis regarding the molecular basis of association signals.

Although relatively few variants are expected to act exclusively through disruption of 3D chromatin contacts, the modular design of the framework permits substitution of the 3D-folding predictor with alternative sequence-to-omics models to evaluate other mechanisms of action. For instance, expression models^61–65^ quantify promoter and enhancer perturbations; splicing models^66^ capture junctional disruptions; chromatin accessibility and transcription factor occupancy models^67,68^ assess regulatory DNA activity; and 3D-architecture models^69^ predict alterations to chromatin contact maps. More comprehensive unified models^70^ aim to integrate these modalities into holistic predictions of regulatory function. In all cases, variant- or haplotype-level in silico mutagenesis yields modality-specific effect scores that can be systematically integrated with GWAS summary statistics, enabling fine-mapping by explicit molecular mechanism.

The *LCOR* gene emerged as a strong candidate in our study with the most significant 3D-genome divergence based on a SNP in its promoter region. *LCOR*, or the *Ligand Dependent Nuclear Receptor Corepressor* gene, is involved in transcriptional repression and chromatin remodeling, suggesting that disruptions in its regulation could impact growth-related pathways in many cells and contribute to height variability^51^. Several other genes identified in this study such as *CCR3*^71^ and *SLC41A2*^72^ also play vital roles in cellular processes, gene regulation, or development, highlighting potential mechanisms through which genetic variation can influence height via 3D-genome architecture.

While this study provides insights into gene regulatory mechanisms potentially underlying height-associated loci, there are several limitations to the approach. First, the height GWAS results referenced primarily cover common genetic variants, excluding rarer variants that may exert stronger effects^73^, including those potentially capable of disturbing 3D genome architecture. As a result, some rare but significant signals contributing to 3D genome disturbance may have been missed. Second, we only evaluate the effects of variants on 3D-genome folding and height. Other genetic, epigenetic, and proteomic factors likely contribute more substantially to height variation. Additionally, while the machine learning model used in this study has been shown to perform well within and across cell types^74^, there are some limits to its scope. Most notably, its predictions are based on a small number of cell types, and it may miss cell-type-specific effects. Akita was also not trained on cell types directly relevant to skeletal growth or chondrogenesis, which are central to height regulation^75^, and it does not currently distinguish functional differences across cell types with high specificity. Furthermore, Akita also analyzes interactions within 1 Mb windows, so longer-range chromatin interactions beyond this distance may not be captured^76,77^. Lastly, the focus on height as a model trait was due to the comprehensive mapping of its genetic architecture; however, given differences in the underlying biology and genetic architecture of complex traits, results from height may not apply to other traits. Thus, it will be valuable to survey traits with large-scale GWAS more broadly using our approach.

In conclusion, this study comprehensively analyzed the 3D-genome disturbance of height-associated loci, identifying 107 regions with suggestive evidence of divergence, and 17 regions with large divergence. Most of these differences in contact are predicted to result from single variants. However, a handful of loci had multiple variants on haplotypes that contributed collectively to 3D-genome changes. These findings underscore the value of combining machine learning models with genomic data to evaluate naturally occurring sequences, propose functional hypotheses about causal variants, and provide new insights into the complex relationship between 3D-genome architecture and trait variation. Future studies could build on this framework by incorporating experimental validation using CRISPR-based perturbations in relevant cell types or by applying complementary sequence-based models to assess effects on gene expression, chromatin accessibility, or splicing. Expanding the predictive framework to additional tissues and developmental stages may also improve the interpretability of trait-associated variants across broader biological contexts.

## Data and Code Availability

All analysis scripts supporting this manuscript are available at: https://github.com/BaranziniLab/ML3DgenomeHeight. Additional materials and information are available from the corresponding author upon reasonable request.

## Acknowledgements

We thank the TOPMed Genetics Consortium, the GIANT Consortium for height, and all study participants for providing the invaluable data that made this analysis possible. We are grateful to the Pollard Lab at UCSF and the Gladstone Institutes for their support in applying the 3D genome prediction machine learning model. We also thank Dr. Colin Brand from the Capra Lab at UCSF for providing the prototype of the data visualization script used in this study.

## Disclosures

RMS reports a service contract with Travere.

## Conflict of interest

The authors of the manuscript declare no conflict of interest.

---

**Supplementary figure S1.**
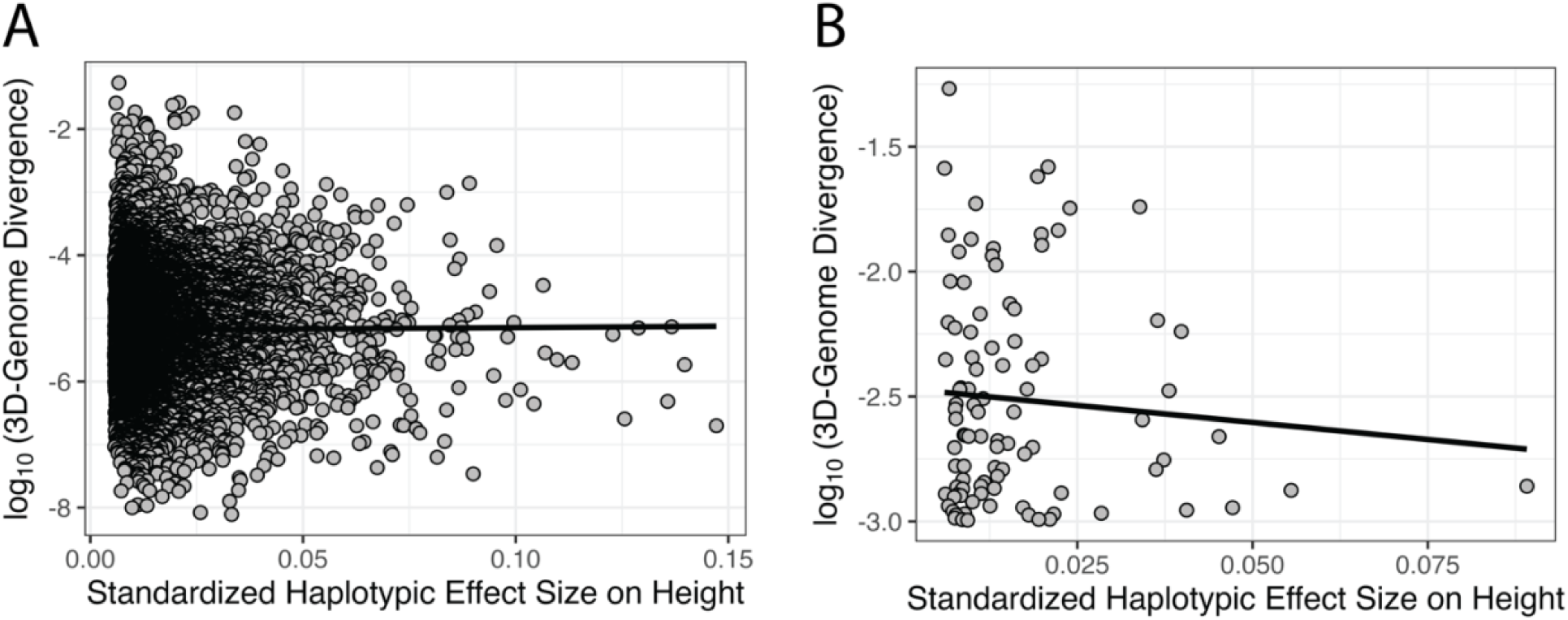
Correlation between haplotypic effect size on height and 3D genome divergence. (A) Scatterplot showing the relationship between the log-transformed maximum predicted 3D genome divergence and the standardized haplotypic effect size on height for all height-associated haplotypes. The linear regression line is overlaid (β = 0.44, p > 0.527), indicating no significant correlation. (B) Scatterplot of the same relationship for height-associated haplotypes in the top 1% for 3D genome divergence. The linear regression line is overlaid (β = −2.73, p > 0.423), again indicating no significant correlation. These results suggest that genetic loci with larger effects on height are not associated with greater 3D genome divergence. Data points represent individual haplotypes.

**Supplementary table s1.** Partitioned heritability of body height within predicted 3D-genome divergent regions. Results from LD Score Regression (LDSC) are shown for African (AFR), Admixed American (AMR), East Asian (EAS), European (EUR), and meta-analyzed (META) GWAS summary statistics. In these analyses, approximately 6.61% of SNPs included in LDSC were annotated as 3D-genome-altering. Columns report the proportion of SNP heritability explained by the annotation (Prop. h²), the estimated enrichment with 95% confidence intervals, and the associated p-values. Across all cohorts and the meta-analysis, enrichment estimates were close to unity and not statistically significant, indicating no evidence for concentration of height heritability within the 3D-genome-altering regions.

| Cohort | Prop. $h^2$ | Enrichment (95% CI) | P-value |
| --- | --- | --- | --- |
| AFR | 0.0831 | 1.257 (0.75, 1.764) | 0.372 |
| AMR | 0.0872 | 1.319 (0.868, 1.769) | 0.151 |
| EAS | 0.0676 | 1.023 (0.829, 1.217) | 0.82 |
| EUR | 0.0947 | 1.431 (0.219, 2.644) | 0.486 |
| META | 0.1006 | 1.521 (0.474, 2.569) | 0.336 |

## Notes

### Competing Interest Statement

The authors have declared no competing interest.

