## Supplementary figures and images for "Evaluating the contribution of genome 3D folding to variation in human height using machine learning"

### Supplementary Figure 1

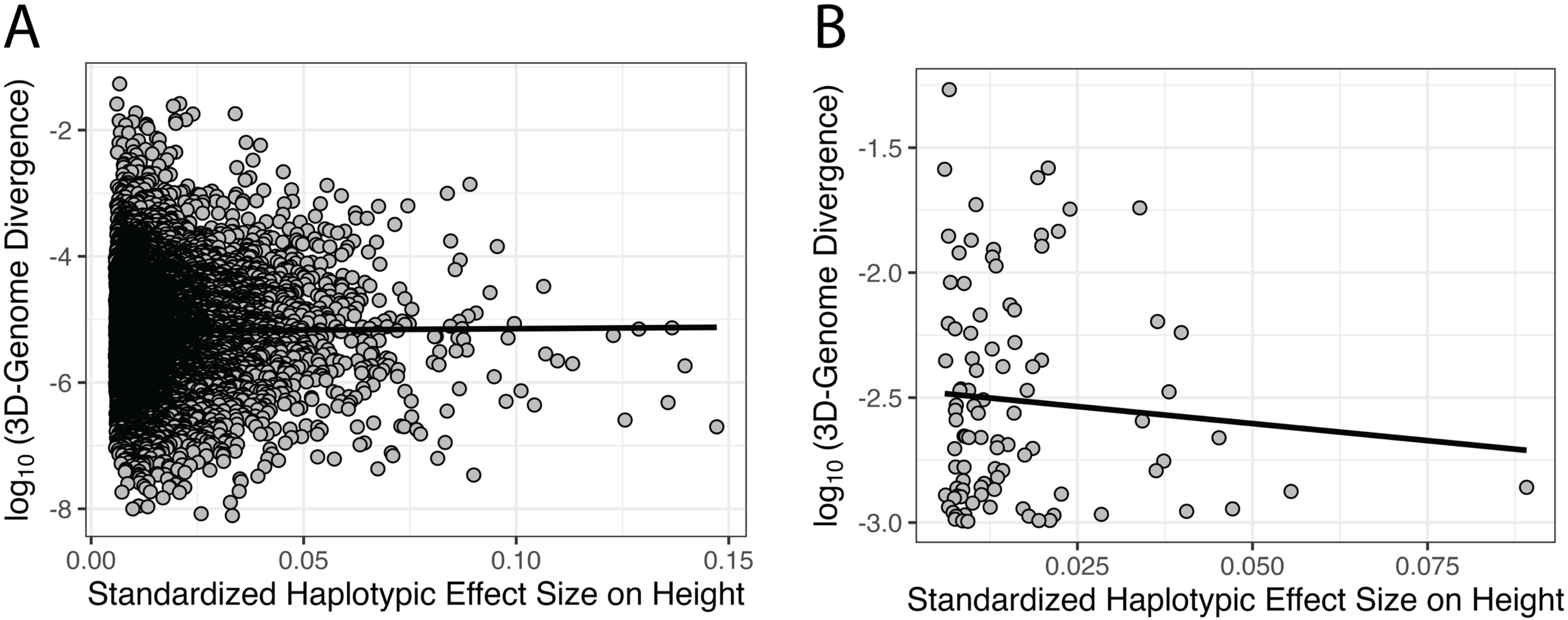
